# Ecological and social drivers of neighbor recognition and the dear enemy effect in a poison frog

**DOI:** 10.1101/2020.04.10.036269

**Authors:** James P. Tumulty, Mark A. Bee

**Author notes:** Department of Neurobiology and Behavior, Cornell University, Ithaca, NY, USA.

## Abstract

Navigating social relationships frequently rests on the ability to recognize familiar individuals using phenotypic characteristics. Across diverse taxa, animals vary in their capacities for social recognition but the ecological and social sources of selection for recognition are often unclear. In a comparative study of two closely related species of poison frogs, we identified a species difference in social recognition of territory neighbors and investigated potential sources of selection underlying this difference. In response to acoustic playbacks, male golden rocket frogs (*Anomaloglossus beebei*) recognized the calls of neighbors and displayed a “dear enemy effect” by responding less aggressively to neighbors’ calls than strangers’ calls. In contrast, male Kai rocket frogs (*Anomaloglossus kaiei*) were equally aggressive to the calls of neighbors and strangers. This species difference in behavior was associated with key differences in reproductive ecology and characteristics of territories. Golden rocket frogs defended reproductive resources in the form of bromeliads, which is expected to create a threat asymmetry between neighbors and strangers favoring decreased aggression to neighbors. In contrast, Kai rocket frogs did not defend reproductive resources. Further, compared with Kai rocket frog territories, golden rocket frog territories occurred at higher densities and were defended for longer periods of time, creating a more complex social environment with more opportunities for repeated but unnecessary aggression between neighbors, which should favor the ability to recognize and exhibit less aggression towards neighbors. These results suggest that differences in reproductive ecology can drive changes in social structure that select for social recognition.

## INTRODUCTION

Social recognition allows individuals to be treated differently according to past interactions and established relationships (Tibbetts and Dale 2007; Wiley 2013a). It underlies a diversity of important social behaviors, such as feeding offspring (Beecher et al. 1981), maintaining pair bonds (Miller 1979), respecting dominance hierarchies (Cheney and Seyfarth 1999), and defending territories (Jaeger 1981). The evolution of social recognition depends on both the benefits of recognition, such as improved mating success or receiving fewer aggressive attacks (e.g., Beletsky and Orians 1989; Sheehan and Tibbetts 2009), as well as the costs associated with the signaling, perceptual, and cognitive mechanisms that make recognition possible (Wiley 2013a). There are now several known cases in which species within a clade differ in the degree to which they exhibit social recognition within the same behavioral contexts, such as nestmate recognition in wasps (Sheehan and Tibbetts 2010), parent-offspring recognition in birds (Beecher et al. 1981; Medvin and Beecher 1986), and territorial neighbor recognition in frogs (Bee et al. 2016). A current challenge to understanding patterns of diversity in recognition systems is identifying sources of selection that favor its evolution in some species but not others (Beecher 1991; Wiley 2013a; Tumulty and Sheehan 2020). Overcoming this challenge requires joint consideration of both ecological and social factors, because many relevant characteristics of social environments depend strongly on underlying variation in ecological factors, such as resource availability (e.g., Hatchwell and Komdeur 2000; Gaulin et al. 2018). Thus far, progress toward identifying ecological and social factors that favor the evolution of social recognition has been limited, in part, due to the necessity of investigating these factors in natural populations of closely related species exhibiting key differences in social recognition.

The recognition of territory neighbors is one of the most widespread forms of social recognition among animals, and it holds considerable promise as a behavioral context for elucidating ecological and social drivers of diversity in social recognition. Evidence for neighbor recognition typically comes from observing the “dear enemy effect” (Wilson 1975; Tumulty 2018a), which occurs when territory holders are less aggressive to neighbors than to strangers. The dear enemy effect can reduce the costs of unnecessary aggression between neighbors with established territory boundaries (Getty 1987; Beletsky and Orians 1989; Qualls and Jaeger 1991; Temeles 1994). Importantly, there is diversity in this behavior among territorial species, as not all territorial animals recognize their neighbors and treat them as “dear enemies” (Temeles 1994). At present, why neighbor recognition and the dear enemy effect evolve in some territorial species but not others remains uncertain. Theoretical considerations, however, identify three key ecological and social factors expected to determine the adaptive value of neighbor recognition and the dear enemy effect, and indeed that of social recognition more broadly.

First, the fitness payoffs resulting from interactions with members of different social categories, such as “neighbor” versus “stranger” (or “dominant” versus “subordinate” or “offspring” vs. “unrelated individual”), determine the adaptive value of behaviorally discriminating between such categories (Getty 1987; Reeve 1989; Sherman et al. 1997; Wiley 2013b). In the context of territorial behavior, the “relative threat hypothesis” (Getty 1987; Temeles 1994) holds that exhibiting lower levels of aggression towards neighbors than strangers is adaptive in situations where neighbors pose less of a threat to territory holders than strangers. The defense of limited resources that are critical for reproduction can produce a threat asymmetry in which neighbors, who already possess resources of their own, are less likely to usurp or make use of the resources of their neighbor than are strangers, who may represent “floaters” without resources (Getty 1987; Temeles 1994). Thus, neighbor recognition and the dear enemy effect are predicted to occur in species whose reproductive ecology involves the defense of limited reproductive resources compared with related species that do not defend such resources.

Second, social recognition should evolve when the complexity of the social environment renders simpler mechanisms of behavioral discrimination (e.g., “rules of thumb”) inadequate. More complex social environments are those in which individuals have more social partners and interact more frequently with those social partners (Freeberg et al. 2012; Bergman and Beehner 2015). Such environments place greater demands on recognizing individuals and managing social relationships (Freeberg et al. 2012). In territorial species, the spatial density of territories is a key aspect of social complexity that should both impact the adaptive value of neighbor recognition and the dear enemy effect and depend on a variety of ecological factors, such as resource abundance and distribution. Territories that occur at higher spatial densities potentially create more opportunities for repeated aggressive interactions with both neighbors and strangers. Avoidance of repeated aggressive interactions with recognized neighbors underlies the benefits of the dear enemy effect. Therefore, neighbor recognition and the dear enemy effect are predicted to occur in species that defend territories at higher densities compared with related species that defend more widely dispersed territories.

Finally, the net benefits of social recognition accrue through time (Getty 1987; Reeve 1989), such that longer social relationships should increase the adaptive value of recognition. Among territorial species, defending territories for longer periods of time potentially provides more opportunities for repeated, but potentially costly and unnecessary, aggressive interactions between the same neighbors. Thus, neighbor recognition and the dear enemy effect are predicted to occur in species that exhibit longer territory occupancies compared with related species that defend territories for shorter periods of time.

Previous research on territorial frogs has revealed taxonomic diversity in neighbor recognition and the dear enemy effect among distantly related species (Bee et al. 2016); however, diversity among closely related species, and how that diversity maps onto species differences in various ecological and social factors, have not been investigated. The variation among frog species in whether neighbors are recognized and treated as dear enemies, coupled with the diversity of territorial behavior in frogs (Wells 1977; Wells 2007; Bee et al. 2013), makes them a potentially informative taxonomic group for comparative research on social recognition. Here, we studied two species of territorial poison frogs – golden rocket frogs (*Anomaloglossus beebei*; Aromabatidae) and Kai rocket frogs (*Anomaloglossus kaiei*) (Fig. 1a) – to investigate potential ecological and social sources of selection favoring the evolution of neighbor recognition and the dear enemy effect. These two congeneric poison frogs are closely related (Grant et al. 2017; Vacher et al. 2017) and last shared a common ancestor some 6 to 12 million years ago (Kumar et al. 2017). Among poison frogs (families Dendrobatidae and Aromobatidae), variation in reproductive ecology is closely linked with variation in social structures and social behaviors, such as territoriality, mating systems, and parental care (Summers and Tumulty 2013). Observations made by us and others (Bourne et al. 2001; Kok et al. 2006; Pettitt et al. 2018; Pettitt et al. 2019) indicate that golden rocket frogs and Kai rocket frogs differ in a key aspect of reproductive ecology that is expected to impact the types of territories that males defend and their relationships with neighbors. Specifically, golden rocket frogs are phytotelm breeders, living and breeding in large terrestrial bromeliads (*Brocchinia micrantha*), where they deposit eggs and tadpoles in small pools of water that collect in leaf axils (“phytotelmata”). Males care for egg clutches in their territories and transport tadpoles on their backs between phytotelmata. In contrast, Kai rocket frogs are terrestrial breeders like most other members of Aromobatidae. Males call from the forest floor and transport tadpoles from terrestrial oviposition sites to deposition sites in pools of water on the forest floor or in bromeliad phytotelmata. Terrestrial breeding, as exhibited by Kai rocket frogs, is ancestral in Aromobatidae (Summers and Tumulty 2013), indicating that phytotelm breeding and any associated behaviors are evolutionarily derived in golden rocket frogs. Males of both species advertise territory ownership vocally (Fig. 1b) and respond aggressively to nearby calling males. Aggressive responses often escalate from aggressive calls (Fig. 1c) to phonotaxis towards the intruder and, eventually, to physical attacks consisting of wrestling and chasing. Thus, because these two species are closely related, similarities in their vocal and aggressive behaviors coupled with differences in their reproductive ecology and territoriality provide a useful comparative system in which to investigate neighbor recognition and the dear enemy effect.

**Figure 1.**
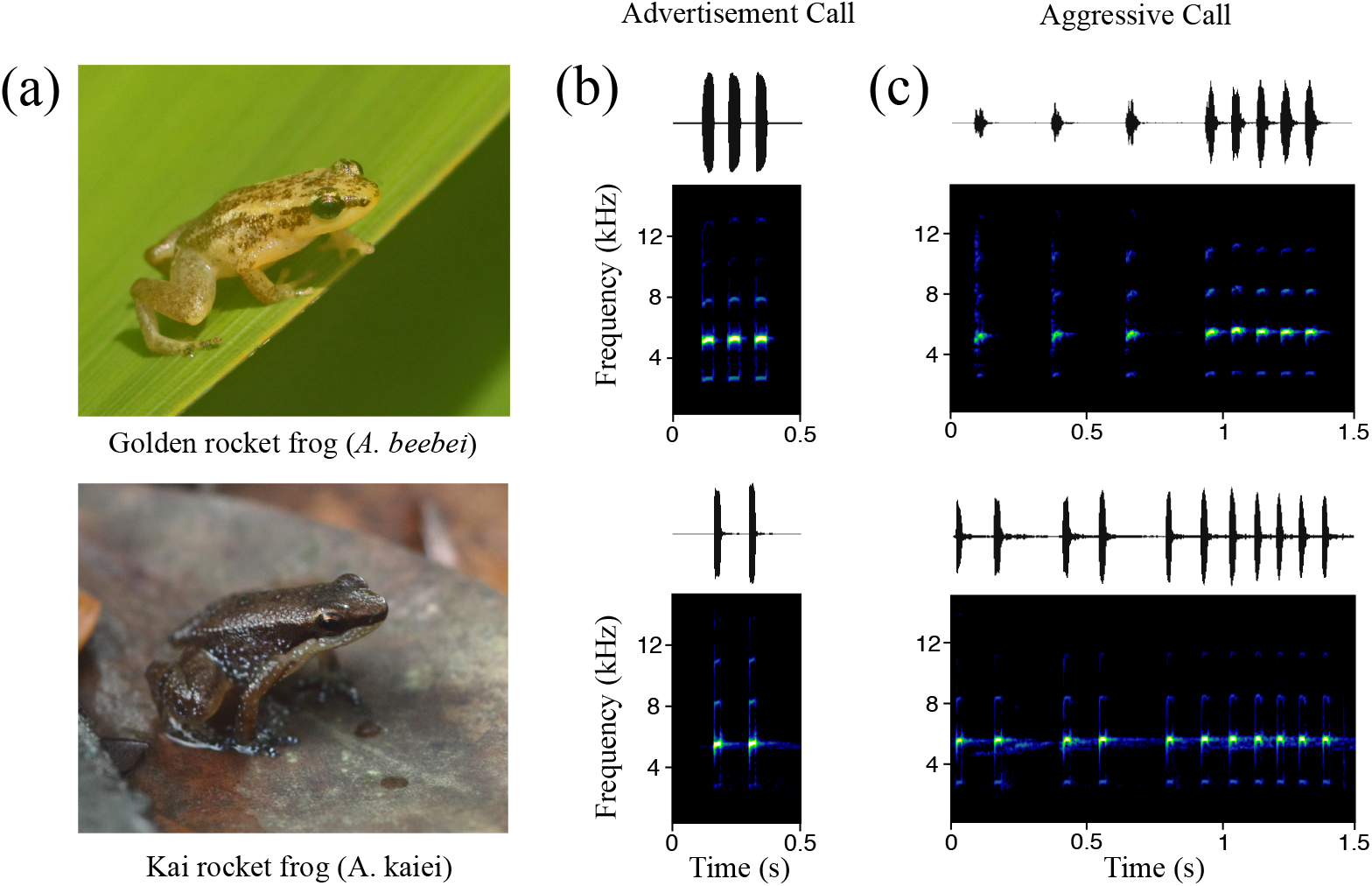
Golden rocket frogs and Kai rocket frogs are close relatives with similar vocal and aggressive behaviors but different reproductive ecologies. (a, top) Male golden rocket frog on a bromeliad leaf and (a, bottom) male Kai rocket frog on the forest floor. (b, c) Waveforms (amplitude over time) and spectrograms (frequency over time) of typical (b) advertisement calls and (c) aggressive calls of (top) golden rocket frogs and (bottom) Kai rocket frogs.

This study had two objectives. First, we tested for neighbor recognition and the dear enemy effect in both species with a field playback experiment to determine whether males recognize the calls of neighbors and respond less aggressively to the calls of neighbors than the calls of strangers. We simulated territory intrusion by broadcasting advertisement calls of neighbors and strangers at increasing amplitudes to determine the territory holder’s aggressive threshold (Rose and Brenowitz 1991), which we defined as the highest stimulus amplitude that a male tolerated without responding aggressively. Higher aggressive thresholds (i.e., lower aggression) to the calls of neighbors than to the calls of strangers constitutes evidence of neighbor recognition and the dear enemy effect (Chuang et al. 2017; Tumulty et al. 2018). Second, we investigated potential sources of selection for neighbor recognition and the dear enemy effect by examining the abovementioned factors predicted to influence the adaptive value of neighbor recognition and the dear enemy effect: (1) the types of resources that are defended, (2) the spatial density of territories, and (3) the duration of territory ownership. To quantify these territory characteristics, we conducted an intensive, multi-year, mark-recapture study to map the territories of males in relation to each other and to the spatial distribution of reproductive resources and to estimate the duration of territory occupancy. To determine the types of resources that are defended we investigated two key criteria of reproductive resource defense (Poelman and Dicke 2008): territories are spatially associated with reproductive resources and offspring (eggs and tadpoles) are located within territories.

## METHODS

### Study site and species

We studied golden and Kai rocket frogs in Kaieteur National Park, Guyana, for five consecutive field seasons from 2013 to 2017. Both species are common in the park. Golden rocket frogs breed year-round in the giant tank bromeliads (*Brocchinia micrantha*) that grow along the plateau near Kaieteur Falls (Bourne et al. 2001; Kok and Kalamandeen 2008; Pettitt et al. 2018; Pettitt et al. 2019). Breeding activity in golden rocket frogs is highest during the rainy season from May-July. Kai rocket frogs are terrestrial frogs that are common in the leaf litter of forested habitats in the park (Kok et al. 2006; Kok and Kalamandeen 2008). Calling by male Kai rocket frogs was observed in all months of our study (April-August), but activity was highest in March and April, the months preceding the onset of the rainy season. Males of both species produce conspicuous advertisement calls (Fig. 1b). These calls consist of series of 3 pulses (range 1-6) for golden rocket frogs or 2 pulses (range 1-2) for Kai rocket frogs with dominant frequencies in the range of 4.6-5.8 kHz (Bourne et al. 2001; Kok et al. 2006; Kok and Kalamandeen 2008; Pettitt et al. 2012; Pettitt et al. 2013). Aggressive calls are longer and consist of several introductory pulses (in golden rocket frogs) or rapid advertisement calls (in Kai rocket frogs) followed by a long train of pulses with relatively shorter inter-pulse intervals (Pettitt et al. 2012; unpublished data; Fig. 2c). Because of variation in the presence of introductory pulses produced by golden rocket frogs, we classified all calls with at least seven pulses as aggressive calls in this species. We monitored individual frogs through time by marking them with unique toe clips and taking dorsal and side-profile photographs of each individual. A combination of toe clips and photographs allowed for identification when frogs were re-captured within and across years.

**Figure 2.**
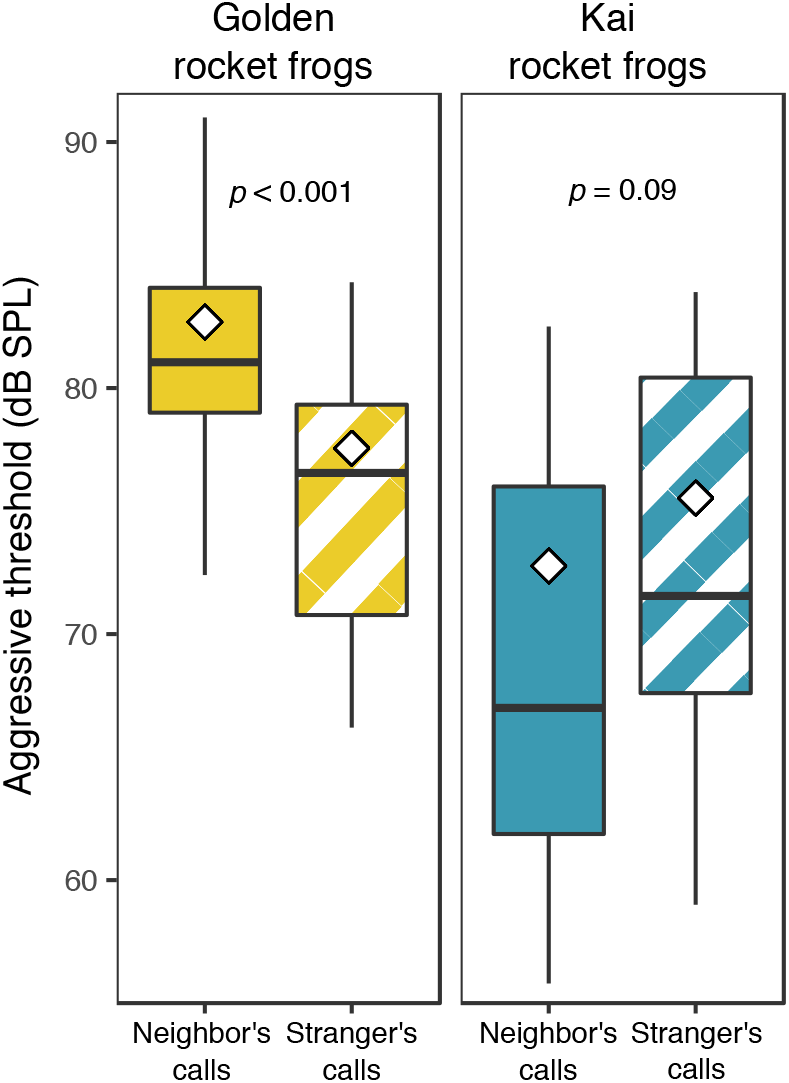
Aggressive thresholds of male golden rocket frogs and Kai rocket frogs to playbacks of the calls of neighbors and strangers. An aggressive threshold is the highest stimulus amplitude a male tolerated before responding aggressively with aggressive calls or approach movements. Black horizontal lines within boxes represent medians and white diamonds represent means. Means were calculated from the absolute sound pressure levels in Pa and re-converted to the dB scale. Box hinges indicate 75th and 25th percentiles, and whiskers extend to the range but no farther than 1.5 times the interquartile range.

### Playback stimuli

To generate stimuli for neighbor-stranger playbacks, we recorded at least two continuous minutes of advertisement calling by previously identified territorial males from a distance of 0.5-1 m using a microphone (Sennheiser ME-66, Wedemark, Germany) and digital recorder (Marantz PMD-620, Kanagawa, Japan; 44.1 kHz sampling rate, 16-bit resolution). After recording, we caught the frog to confirm its identity and measured the air temperature at the site of calling. Although air temperature can affect acoustic properties of anuran vocalizations (Gerhardt and Huber 2002), there is very little temperature variation during the times when these species regularly call. The ranges of temperatures at which recordings were made were 22.8 to 25.8°C for golden rocket frogs and 21.2 to 25.2°C for Kai rocket frogs. Over these ranges there is very little temperature-dependent variation in acoustic properties of calls (Pettitt et al. 2013).

Neighbor and stranger stimuli were created by editing recordings of advertisement calls of specific individuals using Adobe Audition 1.5 (Adobe Systems Inc., San Jose, CA, USA). We standardized the call rate to species-specific means (1 call/2.5 s for golden rocket frogs and 1 call/s for Kai rocket frogs; Pettitt et al. 2012, unpublished data) to create 1-minute stimuli consisting of 23 unique calls for golden rocket frogs or 59 unique calls for Kai rocket frogs by inserting an appropriate amount of silence between consecutive calls. Thus, our stimuli incorporated the natural within-individual variation in calls that would occur over a 1-minute period of calling. Stimuli were high-pass filtered above 2 kHz to minimize background noise, normalized to 80% of their maximum amplitude, and saved as.WAV files (44.1 kHz sampling rate, 16-bit resolution). We then created a series of attenuation levels of each stimulus that differed by 2-dB steps, which allowed us to gradually increase the amplitude of the stimulus in the field. For golden rocket frogs, we created 7 attenuation levels ranging from −12 dB to −0 dB. For Kai rocket frogs, we created 11 attenuation levels ranging from −20 dB to −0 dB because preliminary field playbacks revealed that males of this species sometimes respond aggressively to very low-amplitude playbacks.

### Neighbor-stranger playbacks

We conducted playbacks on males with established territories and previously identified and recorded neighbors. Males were considered territorial if they were observed calling regularly and were captured on at least three different days within an area of typical territory size (Tumulty 2018b). We preferentially tested males with nearby neighbors to maximize the likelihood that they were familiar with their neighbors’ calls. We played back the calls of neighbors and strangers to subjects on the same day, so each subject experienced two playback tests. The stimulus order (whether a neighbor or stranger stimulus was presented first) was randomly assigned to each subject and balanced to result in equal sample sizes for the two possible stimulus orders. We used calls from a subject’s nearest neighbor as the neighbor stimulus. For stranger stimuli, we used calls recorded from males at a different site (see below). All sites were separated by at least 50 m. We replicated stranger stimuli such that each subject heard a different stranger stimulus, except for two stimuli that were each used twice because of limited stimuli available at the times of those tests. Neighbor stimuli were only used more than once for cases in which subjects shared the same nearest neighbor.

Playbacks were performed on actively calling males during times of peak calling activity for both species (06:00 – 11:00), resulting in temperature ranges for playbacks (22.2 to 25.6°C for golden rocket frogs and 21.6 to 24.6°C for Kai rocket frogs) that were very similar to those for acoustic recordings (see above). Stimuli were played to males in the field from a digital audio player (iPod, Apple, Cupertino, CA, USA) connected to an amplified field speaker (Saul Mineroff Electronics, Elmont, NY, USA). The speaker was flat (+/- 2.4 dB) across the dominant frequency range (4.6-5.8 kHz) of both species’ calls. We calibrated the speaker so that the unattenuated stimuli (i.e., −0 dB) were produced at sound pressure levels (SPL) of 82 dB at 1 m, which is at the high end of the range of natural variation in call amplitude for both species (Pettitt et al. 2012; unpublished data). We positioned the speaker facing the subject, along an axis between the subject and its neighbor, at a distance from the subject of 1 m for golden rocket frogs or 1.5 m for Kai rocket frogs. We used different speaker distances for the two species because of natural differences in inter-male distances between the species (Tumulty 2018b). For golden rocket frogs, which called exclusively from bromeliads, the speaker was placed on a tripod at equivalent height to that of the subject and positioned so that it contacted a bromeliad leaf to simulate a realistic calling location. For Kai rocket frogs, the speaker was placed on the ground. If the neighbor was calling at the time, it was either caught and held in a small plastic container for the duration of the test or an observer was positioned close enough to the neighbor to disrupt any calling during the test. Captured neighbors were released immediately after the playback test was completed. Subjects were sometimes disturbed and ceased calling during speaker set-up, so we waited 10 min or until the male started calling again to begin the playback. Nine out of 22 golden rocket frogs and 14 out of 20 Kai rocket frogs were calling at the beginning of the first test. An observer sat quietly 1-2 m away from the subject to observe behavior. Acoustic responses of subjects were recorded using the microphone and digital recorder.

We began playbacks with the lowest amplitude of the stimulus and increased the amplitude in 2-dB steps with each repetition of the 1-min stimulus until a subject responded aggressively or until the highest stimulus amplitude (i.e., −0 dB attenuation) was reached. An aggressive response was noted if the subject either produced an aggressive call or exhibited phonotaxis of at least 20 cm from their original location towards the speaker. We chose 20 cm as our criterion to exclude shorter repositioning movements that males sometimes make while calling. After the first test ended, we waited at least 10 min before beginning the second test, during which time subjects often resumed normal activity. Six out of 22 golden rocket frogs and nine out of 20 Kai rocket frogs were calling at the start of the second test. If the subject had moved in response to the first stimulus, and if it did not return to its original position during the 10-min break, we moved the speaker so that it was again at an appropriate distance and waited an additional 10 min to begin the second test. After the second test, we captured the subject to confirm its identity. We measured the amplitude (dB SPL re 20 *μ*Pa, fast RMS, A-weighted) of the lowest stimulus attenuation level that the subject tolerated without producing an aggressive response at the subject’s location using a digital sound level meter (407764, Extech, Waltham, MA, USA). For subjects that did not respond aggressively, we measured the amplitude of the lowest possible attenuation level (i.e., −0 dB) as a conservative estimate of their aggressive threshold.

We required that subjects respond aggressively to at least one of the two stimuli they heard to ensure they were motivated to respond aggressively and that the range of amplitudes to which they were exposed was sufficiently high to elicit aggression. We excluded trials in which subjects responded aggressively to the initial (quietest) attenuation level of either stimulus, which we did not consider to be a threshold (*n* = 4 trials on 3 individuals for golden rocket frogs and *n* = 1 trial on 1 individual for Kai rocket frogs). Subjects that did not meet these criteria were re-tested on different days. Our final sample sizes were 22 male golden rocket frogs and 20 male Kai rocket frogs. Data were analyzed using the ‘lme4’ package (Bates et al. 2015) in R (R Core Team 2017). We modeled aggressive thresholds using a linear mixed effect model and we modeled whether or not subjects responded aggressively to stimuli using a logistic mixed effects model. We included stimulus (neighbor or stranger), species, and their interaction as fixed effects, while controlling for stimulus order as a fixed effect and individual as a random effect. We then fit separate models for each species to directly compare the responses to neighbors’ and strangers’ calls within species. Significance of fixed effects was tested using Wald chi-square tests with a significance criterion of α = 0.05.

### Territory characteristics

We investigated characteristics of territories that are likely to influence the adaptive value of the dear enemy effect using mark-recapture methods to map the home ranges of territorial males in relation to each other and to the spatial distribution of reproductive resources. We monitored the space use of individual frogs at eight sites (two for golden rocket frogs and six for Kai rocket frogs) in areas of high densities of each species. At each site, we set up a grid with reference flags placed every 2.5 m. The size of each site was selected to encompass all of the calling males within a surveyable area, with site areas ranging from 100 to 2,500 m^2^. Monitoring generally occurred in during times of peak reproductive activity for each species (March-May for Kai rocket frogs and May-August for golden rocket frogs). We visited each site several times per week during the morning (06:00-11:00) and attempted to catch every calling male. Frogs were released within approximately five minutes after being either marked (toe-clipped and photographed) or identified if they had been previously marked, and their locations within the grid were recorded by measuring the distance and compass bearing from the nearest reference flag. Two sites, one for each species (golden rocket frogs: 22.5 x 22.5 m, Fig. 3c, Fig. S1; Kai rocket frogs: 55 x 55 m, Fig. 3d, Fig. S2), were surveyed intensively for multiple years to monitor individual frogs over successive years and to compare the distributions of home ranges to the locations of bromeliads, egg clutches, and tadpoles. Bromeliads were mapped by recording the location of the center of each plant as well as the leaf diameter (distance from the tip of one leaf to the tip of a leaf on the opposite side of the plant) and the diameter of the water-holding center of the plant (distance from the outermost edge of water on one side of the plant to the outermost edge of water on the opposite side of the plant). We also recorded the locations of all the egg clutches and tadpoles we found. Tadpoles were easy to locate by shining a light into bromeliad pools. Because parents sometimes move tadpoles between pools, to avoid potentially re-sampling the same tadpoles we recorded the locations of tadpoles once per field season by systematically searching for tadpoles over the course of several consecutive days. Field locations (distance, compass bearing, and reference flag data) were converted into Cartesian coordinates and analyzed in R as spatial data. We restricted our analysis to adult males that had been observed calling and had at least three capture locations during a field season. We computed each male’s home range as the minimum convex polygon encompassing 100% of its capture locations using the ‘chull’ function in the ‘grDevices’ package and the ‘Polygon’ function in the ‘sp’ package (Bivand et al. 2013).

**Figure 3.**
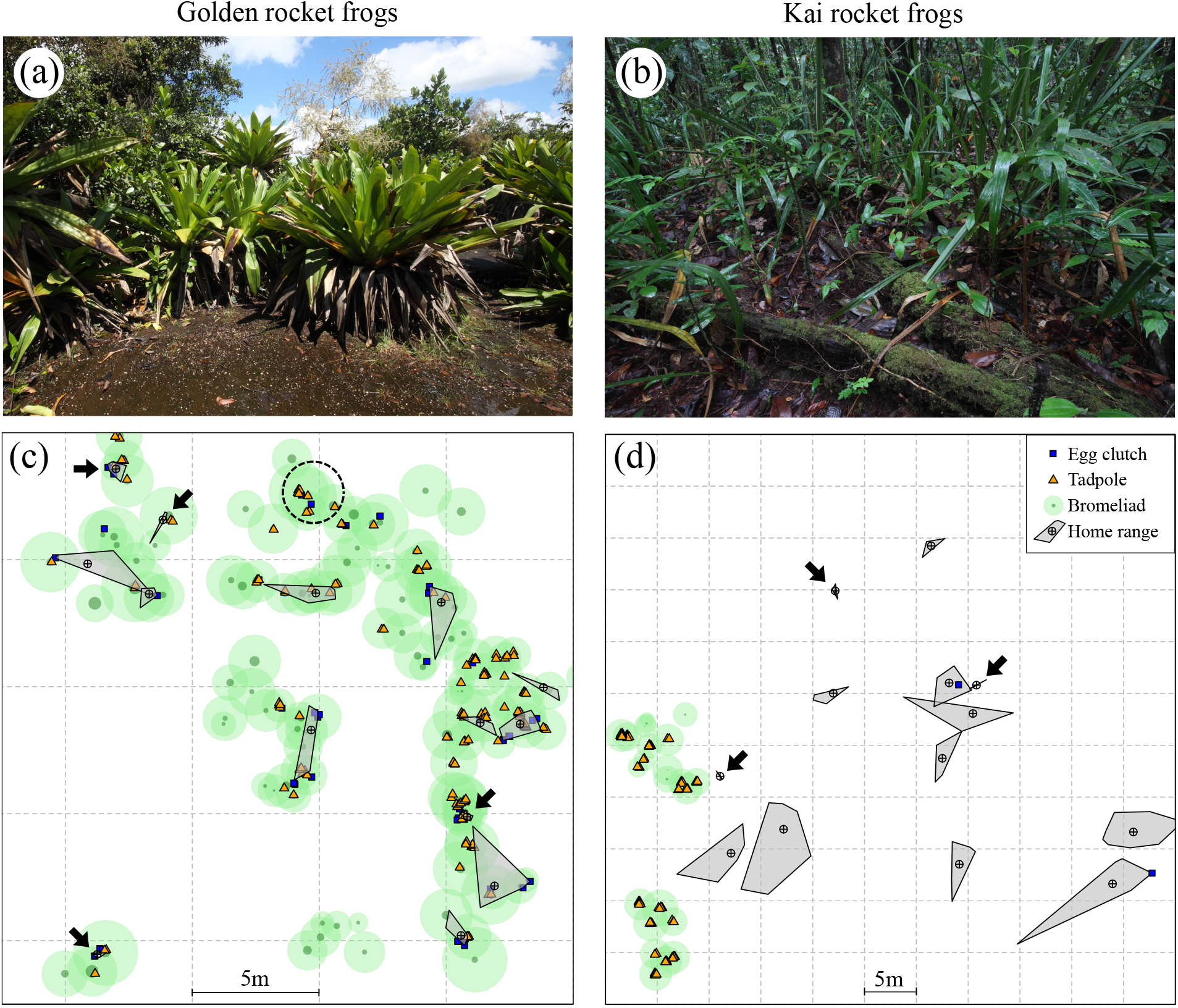
Comparison of habitat use and home range distributions of territorial male golden rocket frogs and Kai rocket frogs. (a, b) Photos of representative habitat at Kaieteur National Park. (a) Habitat of golden rocket frogs consisting of spatially clumped giant tank bromeliads (*Brocchinia micrantha*) with relatively bare rock between clumps. (b) Habitat of Kai rocket frogs consisting of typical rainforest understory. (c, d) The spatial distribution of male home ranges (gray polygons), bromeliads (green circles), eggs (blue rectangles), and tadpoles (orange triangles) at sites surveyed for (c) golden rocket frogs and (d) Kai rocket frogs in 2015 (maps for other years are in the supporting information). Black arrows point to males with particularly small home ranges. Note: these sites are on different scales; the dashed gray lines represent a grid at 5m intervals. The dotted circle in (c) represents an area with a territorial male that we were unable to regularly catch and hence is not represented with a home range. Light green circles represent the leaf diameters of bromeliads, and smaller dark green circles represent the phytotelmata diameters within bromeliads.

To test the prediction that neighbor recognition and the dear enemy effect are more likely to occur in species that defend reproductive resources, we determined whether territories are spatially associated with the distribution of reproductive resources, which is one of the key criteria of reproductive resource defense (Poelman and Dicke 2008). We represented bromeliads based on their measured leaf diameter as spatial polygons using the function ‘gBuffer’ in the ‘rgeos’ package (Bivand and Rundel 2017). Because repeated locations of individual males are not independent, we randomly selected one location per male and computed the proportion of these locations that were within bromeliad polygons at each site. To control for the different availability of bromeliads at the different sites, we used a Monte Carlo approach to compare the observed distributions of males to random locations at each site (Brown et al. 2009), which were simulated using the ‘spsample’ function in the ‘sp’ package (Bivand et al. 2013). Specifically, the observed proportion of *n* capture locations on bromeliads was compared to the distributions of proportions of simulated data on bromeliads from 10,000 iterations of *n* random locations at each site. We then computed p-values as the proportion of simulated proportions that were equal to or more extreme than the observed proportion.

We investigated the second criterion of reproductive resource defense – that eggs and tadpoles are found within territories – by first estimating territory boundaries. To do this, we added a species-specific buffer around each male’s home range, because the area that males defend is likely larger than the area in which they typically call (Ringler et al. 2011). This buffer (0.87 m for golden rocket frogs and 1.75 for Kai rocket frogs) estimated the average distance at which a calling stranger would be tolerated before receiving aggression from a territorial male. We computed these distances from the aggressive thresholds determined in the neighbor-stranger discrimination experiment and models of sound attenuation based on field measurements of the relationship between SPL and distance for each species’ calls in species-specific habitats (unpublished data). These models were *SPL* = 74.57 – 10.57 × *ln*(*meters*) for golden rocket frogs and *SPL* = 77.69 – 11.87 × *ln*(*meters*) for Kai rocket frogs. We then computed the proportions of egg clutches and tadpoles that were within these territories. We again compared the observed proportion of egg clutches and tadpoles within territories to distributions of simulated random points at each site, using the previously described simulation procedure.

To test the prediction that neighbor recognition and the dear enemy effect are more likely to be found in species that defend territories occurring at higher spatial densities, we quantified home range size and nearest neighbor distances for territorial males from all eight sites. Home range size was computed as the area of the minimum convex polygon determined from each male’s capture locations. Nearest neighbor distance was calculated as the Euclidian distance between the centroid of a male’s capture locations and the centroid of its nearest neighbor’s capture locations. Home range data and nearest neighbor distance data were log-transformed to improve linearity for statistical analyses. We tested whether home range size and nearest neighbor distance differed between species using linear mixed effects models implemented through the ‘lme4’ package; the number of capture locations per individual and individual were included as random effects due to the positive relationship between the number of capture points and estimates of home range size and repeated sampling of individuals across years. Significance was evaluated using Wald chi-square tests.

To test the prediction that neighbor recognition and the dear enemy effect are more likely to occur in species that have longer territory occupancies, we estimated of the duration of territory occupancies for the two sites that were monitored intensively over multiple years. For Kai rocket frogs, which had a more discrete breeding season, we report the times between the date of first and last capture for each territorial male. However, we emphasize that these are likely underestimates because some males were already calling and defending territories when we began monitoring in March of each year. For golden rocket frogs, which often maintained the same territories for the entire duration of our field seasons (March-August), we instead report data on the number of years in which males were observed in the same territories.

## RESULTS

### Neighbor-stranger playbacks

Both species responded aggressively to playbacks, but only golden rocket frogs had relatively higher aggressive thresholds to the calls of neighbors. Males of both species typically oriented towards the speaker and produced advertisement calls in response to the initial lower amplitude stimuli. Aggressive responses of golden rocket frogs (*n* = 22) to higher amplitude stimuli typically consisted of aggressive calls (*n* = 25 of 44 tests) and occasionally phonotaxis (*n* = 5 of 44 tests). Kai rocket frogs (*n* = 20) often responded aggressively with phonotaxis (25 of 40 tests) and occasionally with aggressive calls (9 of 40 tests). In a statistical model of aggressive thresholds, there was a significant interaction (χ^2^ = 13.10, p < 0.001) between stimulus (neighbor vs. stranger) and species (golden rocket frogs vs. Kai rocket frogs) (Fig. 2). Stimulus order had no overall effect on aggressive thresholds (χ^2^ = 2.11, *p* = 0.15). The aggressive thresholds of golden rocket frogs were significantly higher, on average by 4.7 dB, for the calls of neighbors than the calls of strangers (χ^2^ = 15.58,*p* < 0.001, Fig. 2). This amplitude difference corresponds approximately to a 36% difference in distance at which calling neighbors would be tolerated (0.56 m) compared with strangers (0.87 m), based on a model of sound attenuation. Golden rocket frogs were also more likely to respond aggressively to the calls of strangers (19 of 22 males) than to the calls of neighbors (9 of 22) (χ^2^ = 8.26, *p* = 0.004). In contrast, for Kai rocket frogs, there was no significant difference in their aggressive thresholds for the calls of neighbors and strangers (χ^2^ = 2.95, *p* = 0.09, Fig. 2) or in their likelihood of responding aggressively to the two stimuli (χ^2^ = 3.22, *p* = 0.07). Based on their overall mean aggressive threshold (71 dB SPL), we estimated that male Kai rocket frogs would tolerate calling conspecific males at an average distance of 1.75 m, slightly more than twice the distance that male golden rocket frogs tolerate conspecifics.

### Territory characteristics

Bromeliads were critical reproductive resources for golden rocket frogs, as all of the egg clutches (*n* = 141) and tadpoles (*n* = 460) we found were deposited in bromeliad phytotelmata (Fig. 3c, Fig. S1). Further, tadpoles were almost always deposited singly, with only 2% of phytotelmata having more than one tadpole. Male golden rocket frogs defended territories that were spatially associated with these plants; the observed proportion of capture locations for territorial males on bromeliads ranged from 0.89 to 1.0 over five years of monitoring and was always greater than chance (all *p* < 0.001, Table 1, Fig. 3c, Fig. 4a). Further, the proportion of offspring within territories across years was also always greater than chance (egg clutches: range 0.66 to 0.95, all *p* < 0.001; tadpoles: range 0.65 to 0.76, all *p* < 0.001; Table 2, Fig. 4b).

**Figure 4.**
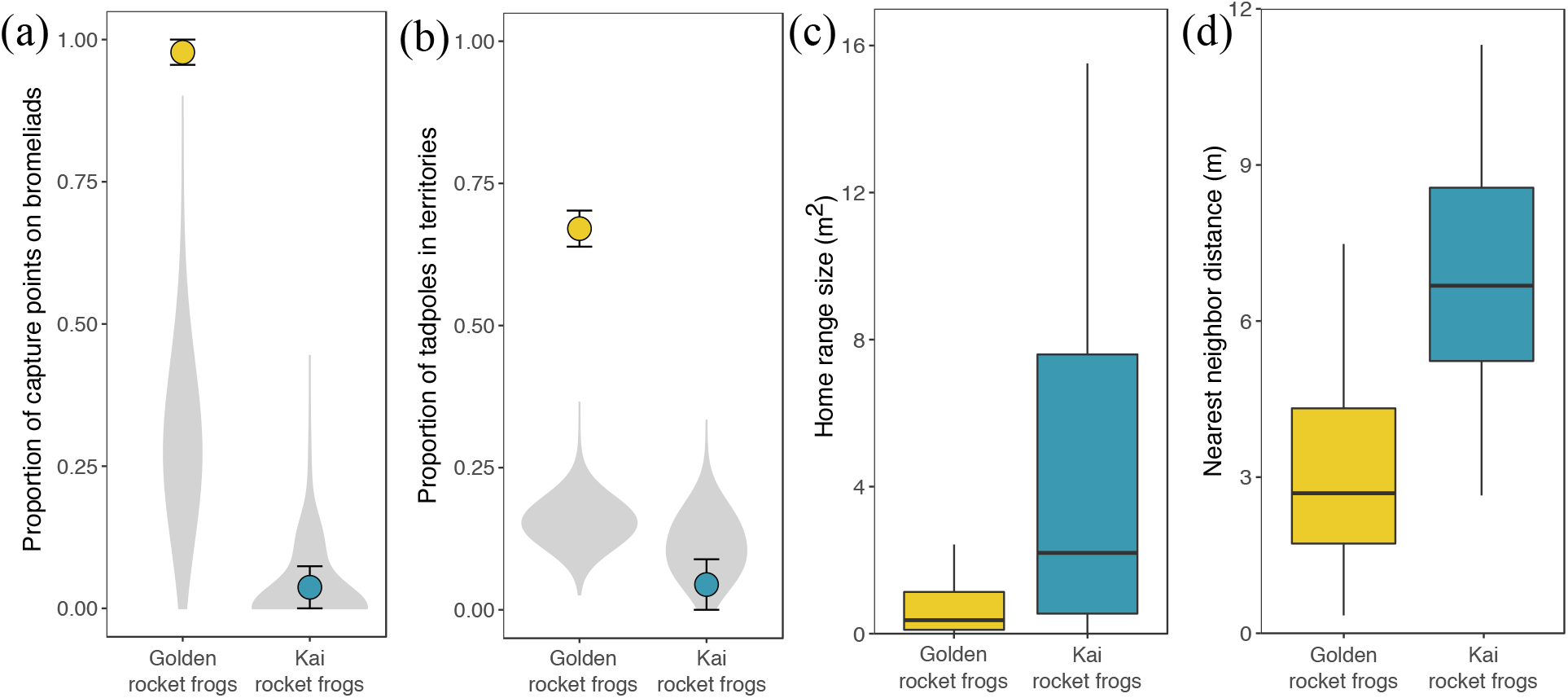
Quantitative species comparisons of home range characteristics. (a, b) Points depict the mean (±SE) proportions of captures points on bromeliads and tadpoles in male territories over multiple years of surveying the same two sites shown in Figure 3. Light gray violin plots represent the distributions of iterations of simulated random locations (a) on bromeliads or (b) in territories, pooled across years. Data for each year are shown in Tables 1 and 2. (c) Home range sizes (100% minimum convex polygons) and (d) nearest neighbor distances of territorial male golden rocket frogs and Kai rocket frogs. Black horizontal lines within boxes represent medians, box hinges indicate 75th and 25th percentiles, and whiskers extend to the range but no farther than 1.5 times the interquartile range.

**Table 1.** Spatial association between the locations of territorial males and bromeliads, compared with randomly distributed locations, at two sites.

| Species | Year | No. males | No. Bromeliads | Observed proportion | Simulated proportion |  |  |
| --- | --- | --- | --- | --- | --- | --- | --- |
|  |  |  |  |  | Mean | (95% CI) | p-value |
| Golden rocket frogs | 2013 | 11 | 93 | 1 | 0.24 | (0, 0.55) | <0.001 |
|  | 2014 | 9 | 128 | 0.89 | 0.28 | (0, 0.56) | <0.001 |
|  | 2015 | 14 | 122 | 1 | 0.31 | (0.07, 0.57) | <0.001 |
|  | 2016 | 10 | 114 | 1 | 0.30 | (0, 0.6) | <0.001 |
|  | 2017 | 15 | 109 | 1 | 0.30 | (0.07, 0.53) | <0.001 |
| Kai rocket frogs | 2015 | 13 | 23 | 0 | 0.03 | (0, 0.15) | 1 |
|  | 2016 | 9 | 24 | 0.11 | 0.03 | (0, 0.22) | 0.52 |
|  | 2017 | 7 | 19 | 0 | 0.03 | (0, 0.14) | 1 |

Observed and simulated proportions represent the proportions of points within the leaf diameter of bromeliads at each site. We randomly selected one location for each male to calculate the observed proportion. Mean and 95% confidence intervals for the simulated proportion were computed from 10,000 iterations of randomly distributed points, each of equal sample size to the number of males at that site-year.

**Table 2.** The observed proportions of tadpoles and egg clutches within male territories compared with the proportions of simulated randomly distributed locations.

| Species | Year | No. tadpoles<br>or clutches | No.<br>males | Observed<br>proportion | Simulated proportion |  |  |
| --- | --- | --- | --- | --- | --- | --- | --- |
|  |  |  |  |  | Mean | (95% CI) | p-value |
| Tadpoles |  |  |  |  |  |  |  |
| Golden<br>rocket frogs | 2014 | 94 | 9 | 0.67 | 0.14 | (0.07, 0.21) | <0.001 |
|  | 2015 | 165 | 14 | 0.76 | 0.18 | (0.12, 0.24) | <0.001 |
|  | 2016 | 74 | 10 | 0.61 | 0.16 | (0.08, 0.24) | <0.001 |
|  | 2017 | 127 | 15 | 0.65 | 0.14 | (0.09, 0.20) | <0.001 |
| Kai rocket<br>frogs | 2015 | 135 | 13 | 0 | 0.16 | (0.10, 0.22) | <0.001 |
|  | 2016 | 45 | 9 | 0.13 | 0.09 | (0.02, 0.18) | 0.49 |
|  | 2017 | 59 | 7 | 0 | 0.08 | (0.02, 0.15) | 0.01 |
| Egg clutches |  |  |  |  |  |  |  |
| Golden rocket<br>frogs | 2014 | 29 | 9 | 0.66 | 0.14 | (0.03, 0.28) | <0.001 |
|  | 2015 | 62 | 14 | 0.85 | 0.18 | (0.10, 0.27) | <0.001 |
|  | 2016 | 31 | 10 | 0.71 | 0.16 | (0.03, 0.29) | <0.001 |
|  | 2017 | 19 | 15 | 0.95 | 0.14 | (0, 0.32) | <0.001 |

Note: we did not find enough Kai rocket frog egg clutches to warrant inclusion in spatial analyses. Simulated proportions are from 10,000 iterations of randomly simulated points, each of equal sample size to the number of tadpoles or egg clutches in a given site-year.

In contrast, male Kai rocket frogs defended territories on the forest floor that were not spatially associated with bromeliads (Fig. 3b, d, Fig. S2). The observed proportion of capture locations of this species on bromeliads ranged from 0.0 to 0.11 over three years of monitoring and never differed from chance (all *p* > 0.05; Table 1, Fig. 4a). Of the three Kai rocket frog egg clutches we found (Fig. 3d), all were deposited within or near male territories under dead leaves on the forest floor. This species did deposit tadpoles (*n* = 239) in bromeliad phytotelmata, which were often located well outside of male territories (Fig. 3d; Fig. S2). The observed proportion of tadpoles in Kai rocket frog territories ranged from 0.0 to 0.13 and was significantly *less than* the proportion expected by chance in 2015 and 2017 (all *p* < 0.05; Table 2, Fig. 4b) and did not differ from chance in 2016 (*p* = 0.49; Table 2, Fig. 4b). Phytotelmata with multiple tadpoles were more common for Kai rocket frogs (22% compared with 2% in golden rocket frogs; χ^2^ = 63.79, *p* < 0.001), and they contained between two and five tadpoles. These results indicate that Kai rocket frogs transport their tadpoles from oviposition sites in the leaf litter within or near their territories to tadpole deposition sites that are outside their territories, where multiple males potentially share the same phytotelm.

The spatial density of territories was much higher for golden rocket frogs than it was for Kai rocket frogs, as golden rocket frogs defended smaller territories and were closer to their neighbors than Kai rocket frogs. Home range sizes of territorial male golden rocket frogs averaged 15.2% of those of Kai rocket frogs (χ^2^ = 27.89, *p* < 0.001, *n* = 108; Fig. 4c) and nearest neighbor distances in golden rocket frogs averaged 42.6% of those in Kai rocket frogs (χ^2^ = 45.73, *p* < 0.001, *n* = 104; Fig. 4d).

Male golden rocket frogs also held territories for longer periods of time than Kai rocket frogs. Within a field season, golden rocket frogs generally occupied the same territories for the entire monitoring period, which was concentrated during the months of May, June, and July. Many of these males also maintained the same territories across field seasons, as approximately half of the golden rocket frogs we monitored each year were recaptures from previous years (Table 3). We identified 10 golden rocket frogs that defended the same territories across 3 years, five that defended them across 4 years, and one male that defended the same territory across all 5 years of the study (Table 3). Kai rocket frogs typically occupied the same territories for several weeks within a field season, from prior to the start of our monitoring period in March until the onset of the rainy season in May, when calling and territorial behavior by Kai rocket frogs decreased substantially. The mean territory duration for Kai rocket frogs was at least 23 days (range = 2 to 46 days). In contrast to golden rocket frogs, Kai rocket frogs were not recaptured between years at a site that we monitored intensively over three years. Anecdotally, at other sites, we observed one male Kai rocket frog that defended the same territory across two years and one that defended the same territory across three years. However, the fact that no multi-year territories were observed at the site we monitored intensively indicates that multi-year territories are rare for male Kai rocket frogs.

**Table 3.** The total numbers of golden rocket frog males encountered each year at a site surveyed for five consecutive years, as well as the number of positively identified recaptured males from previous years.

| Species | Year | No.<br>Males | 2<sup>nd</sup> year<br>recaptures | 3<sup>rd</sup> year<br>recaptures | 4<sup>th</sup> year<br>recaptures | 5<sup>th</sup> year<br>recaptures |
| --- | --- | --- | --- | --- | --- | --- |
| Golden rocket<br>frogs | 2013 | 16 | - | - | - | - |
|  | 2014 | 14 | 4 | - | - | - |
|  | 2015 | 20 | 7 | 4 | - | - |
|  | 2016 | 18 | 2 | 6 | 1 | - |
|  | 2017 | 16 | 3 | 0 | 4 | 1 |

Note: Kai rocket frogs are not included because we did not recapture any Kai rocket frogs between years at a site that was surveyed for three consecutive years. Additionally, the numbers of encountered golden rocket frogs each year are greater than the numbers in Table 1 and 2 because this table includes all males encountered in a year, not just those with at least three capture points.

## DISCUSSION

In this paper we addressed a major question concerning the evolution of social recognition: what are the ecological and social sources of selection that favor recognition? In a study of two closely related poison frogs, we discovered a species difference in neighbor recognition and the dear enemy effect and used this difference to identify likely sources of selection for this common form of social recognition. We found that male golden rocket frogs recognized the calls of their neighbors and responded less aggressively to the calls of neighbors than to the calls of strangers. This result confirms an earlier qualitative report (Bourne et al. 2001). In contrast, male Kai rocket frogs did not behaviorally discriminate between the calls of neighbors and strangers. This species difference in whether neighbors were recognized and treated as dear enemies was associated with a key species difference in reproductive ecology that impacted characteristics of territories and relationships with neighbors. Overall, the species differences we found are consistent with our three predictions that neighbor recognition and the dear enemy effect should be favored in species that defend limited reproductive resources, in territories that occur at higher densities, and in territories that are held for longer periods of time. We propose that the evolutionary transition to phytotelm breeding in golden rocket frogs likely drove these changes in the types of territories that males defend and in the social structure of territories which, in turn, favored the evolution of neighbor recognition and dear enemy behavior in this species.

Several authors have argued that the adaptive value of the dear enemy effect depends on the type of territory that is defended and the nature of relationships between territory holders and their neighbors (Temeles 1994; Müller and Manser 2007; Bee et al. 2016; Christensen and Radford 2018). While studies of changes within species in whether neighbors are treated as dear enemies show that this behavior is indeed dependent on social and ecological context (Hyman 2005; Briefer et al. 2008), large phylogenetic differences between species that do and do not exhibit the dear enemy effect have previously limited the identification of social and ecological sources of selection for this behavior. By comparing two close relatives with otherwise similar behaviors and ecology, we were able to identify three key characteristics of territories that are associated with neighbor recognition and the dear enemy effect in golden rocket frogs.

First, we found that male golden rocket frogs defended reproductive resources, but Kai rocket frogs did not. The defense of limited reproductive resources is expected to create a threat asymmetry between neighbors and strangers (Temeles 1994). When animals defend limited reproductive resources, strangers that they encounter are likely “floaters” who do not possess reproductive resources and may be more likely to attempt a territory takeover than are neighbors, who already have resources of their own (e.g., Booksmythe et al. 2010). In such situations, territory holders should respond to rivals according to the threat that they pose, treating less-threatening neighbors with less aggression. We found that territorial male golden rocket frogs were spatially associated with the distribution of reproductive resources and that eggs and tadpoles were found within male territories, meeting two key criteria of reproductive resource defense (Poelman and Dicke 2008). Bromeliad phytotelmata served as both oviposition and tadpole deposition sites for golden rocket frogs and are therefore critical reproductive resources. Males were nearly always found on bromeliads and defended territories that included one or several of these plants. One alternative interpretation of this spatial association is that males are not defending reproductive resources *per se* but are simply defending territories in suitable habitat, since females and juveniles also live in these bromeliads. However, males care for eggs and tadpoles throughout development, even transporting large, well-developed tadpoles between pools in their territories, suggesting that they are in fact defending phytotelmata that contain their offspring (Bourne et al. 2001; Pettitt et al. 2018; Pettitt et al. 2019). Similar spatial associations between male territories and reproductive host plants have been found for other phytotelm-breeding poison frogs in which males are heavily involved in parental care (Poelman and Dicke 2008; Brown et al. 2009).

In contrast, male Kai rocket frogs defended territories on the forest floor that were not spatially associated with reproductive resources. Eggs were deposited within male territories on dead leaves on the forest floor, which are ubiquitous in rainforests and unlikely to constitute a limited or defended reproductive resource (Donnelly 1989; Roithmair 1992; Pröhl 1997; Ursprung et al. 2011). Males transported tadpoles to bromeliad phytotelmata or terrestrial pools that were outside their territories, where offspring from multiple males may often develop in the same pool (unpublished data). Taken together, we interpret these results as evidence that male Kai rocket frogs do not defend limited reproductive resources. Instead territories in this species likely function as areas where males can advertise to and court females uninterrupted by rivals. Similar interpretations have been made for other poison frogs with similar reproductive ecology (Roithmair 1992; Pröhl 1997; Pröhl and Berke 2001; Ursprung et al. 2011). Thus, one explanation for the lack of neighbor-stranger discrimination in Kai rocket frogs is that neighbors and strangers both represent competitors for mates, and as such, pose equivalent threats to territory owners. In such situations, territory owners should respond with aggression to any rival male encroaching on their territory, regardless of whether they are neighbors or strangers. This explanation has also been proposed for the lack of behavioral discrimination between neighbors and strangers found in territorial male strawberry poison frogs (*Oophaga pumilio*) (Bee 2003) and brilliant-thighed poison frogs (Tumulty et al. 2018), both of which, like Kai rocket frogs, defend territories on the forest floor that are not spatially associated with reproductive resources.

Second, male golden rocket frogs defended smaller territories and were closer to their neighbors than Kai rocket frogs. The spatial density of territories can impact the rates at which territory owners encounter both neighbors and strangers. Higher density territories may limit the efficacy of decision rules based on simpler “rules of thumb” because they represent more complex social environments, with higher interaction rates. Given the costs of aggressive interactions in frogs (Dyson et al. 2013), the ability to recognize neighbors can allow males to cope with the demands of this increased social complexity by minimizing the frequency of aggressive interactions with nearby neighbors but still allowing males to respond aggressively to strangers. The territory sizes and nearest neighbor distances for golden rocket frogs are among the smallest documented among poison frogs (cf. Roithmair 1992; Summers 1992a; Pröhl 1997; Summers 2000; Pröhl and Berke 2001; Poelman and Dicke 2008; Brown et al. 2009; Ringler et al. 2009; Werner et al. 2010; Ringler et al. 2011; Tumulty et al. 2018), suggesting the potential for unusually high rates of aggressive interactions between neighbors. For Kai rocket frogs, which maintain greater inter-male distances, males may be able to rely on a simpler decision rule according to which they respond aggressively to any calling male encroaching on their territory. It will be important in future work to test these ideas directly by quantifying species differences in the rates of aggressive interactions between territory residents and other males.

Finally, golden rocket frogs defended territories for longer durations than Kai rocket frogs. Territories that are held for longer periods of time should also favor neighbor recognition because the net benefits of avoiding repeated but unnecessary aggressive interactions with nearby neighbors accrue over time. The territory tenure of golden rocket frogs was remarkable; it was not uncommon for males to defend the same territory for 3 or 4 consecutive years (Table 3), and we observed one male that maintained the same territory for all 5 years of this study. To our knowledge this is the longest documented territory tenure in a frog (Pröhl 2005; Wells 2007). Such long territory tenure may result from the ability to monopolize stable reproductive resources that often maintain pools of water year round (Bourne et al. 2001). Longer territory tenure could also result in male golden rocket frogs becoming more familiar with their neighbors than is possible for male Kai rocket frogs, and models based on familiarity have been proposed as explanations for the dear enemy effect (Ydenberg et al. 1988; Getty 1989; Temeles 1994). However, we think differences in familiarity with neighbors are unlikely to explain the observed species difference in whether or not territorial males treat their neighbors as dear enemies. This is because even though Kai rocket frogs share territory boundaries with neighbors for shorter durations than golden rocket frogs, they still defend territories for at least several weeks, which is should provide ample opportunity for males to learn their neighbors’ calls. For example, male bullfrogs (Howard 1978) and olive frogs (Chuang et al. 2013) also defend territories for several weeks during the breeding season, and both of these species exhibit neighbor recognition and the dear enemy effect (Davis 1987; Chuang et al. 2017).

This study adds to a growing number of studies in frogs (reviewed in Bee 2016; Bee et al. 2016) and other animals (reviewed in Temeles 1994) demonstrating variation in neighbor recognition and the dear enemy effect among territorial species. Vocally-mediated neighbor recognition has been demonstrated in two frog species in the family Ranidae (Davis 1987; Bee and Gerhardt 2002; Chuang et al. 2017), but studies of some other distantly related frog species, including other poison frogs (Bee 2003; Tumulty et al. 2018), have failed to find robust evidence that territory holders recognize neighbors and behaviorally discriminate between the calls of neighbors and strangers (Bee 2016; Bee et al. 2016). The difference in neighbor recognition and the dear enemy effect between closely related species shown here highlights that this behavior is evolutionarily labile. One potential target of selection for neighbor recognition and the dear enemy effect among frogs is the specificity of aggressive behavioral plasticity. Among lek-breeding territorial frogs, aggressive thresholds for calls are often assumed to be generalized to the calls of conspecific males, such that territory holders respond aggressively to any call that is perceived at an amplitude exceeding the territory holder’s threshold, regardless of whether or not the caller is familiar. These thresholds are also plastic; after exposure to high-amplitude calls, males temporarily elevate their thresholds for responding aggressively to these calls, allowing them to tolerate new calling males that have established territories nearby (Rose and Brenowitz 1991; Marshall et al. 2003; Humfeld et al. 2009; Reichert 2010). Plastic aggressive thresholds thus allow males to modulate their aggression in order to track the local density of calling males. Although we did not specifically investigate plasticity, our results are broadly consistent with such a relationship between density and aggressive thresholds: compared with Kai rocket frogs, golden rocket frogs had overall higher aggressive thresholds that corresponded to a higher density of territorial males. However, the finding that male golden rocket frogs have elevated aggressive thresholds that are specific to the calls of neighbors suggests that selection has acted on this ancestral behavioral plasticity by modifying its specificity. Elevating aggressive thresholds for the calls of individual neighbors based on acoustic properties specific to those individuals while maintaining a lower aggressive threshold for unfamiliar calls would allow males to treat their neighbors as dear enemies (Bee et al. 2016). Future comparative research in frogs is needed to understand the mechanisms underlying the specificity of aggressive plasticity and the extent to which specificity is evolutionarily labile.

In conclusion, our results suggest that the evolutionary transition to phytotelm breeding in golden rocket frogs produced a threat asymmetry between neighbors and strangers and drove changes in social structure, including an increase in social complexity, that together favored the evolution of social recognition of territory neighbors. By defending spatially clumped, stable reproductive resources, the distribution of territorial males was likely constrained to the distribution of those resources, putting males in closer proximity to neighbors and for longer periods of time than they would be if they were not defending such resources. Kai rocket frogs, in contrast, may be able to maintain greater inter-male distances, unconstrained by the distribution of reproductive resources, allowing them to defend territories by using simpler decision rules of responding aggressively to any conspecific male. Additional studies in other species are needed to determine if these factors are general predictors of neighbor recognition and the dear enemy effect both among frogs and among other species. Regardless, reproductive ecology has been identified as a key factor that can influence social structure in frogs (Poelman and Dicke 2008; Brown et al. 2009) and other animals (Ostfeld 1985; Fincke 1992; Rubenstein and Lovette 2007), and our study extends this body of work to the evolution of social recognition that allows animals to navigate changes in social structure.

## Acknowledgments

We sincerely thank Godfrey Bourne and Beth Pettitt for advice and logistical assistance, and Calvin Giddings, Mahendra Doraisami, Maxwell Basil, Thomas John, Andrius Pašukonis, Zachary Lange, Chloe Fouilloux, and Johana Goyes Vallejos for help in the field. We thank Jason Brown and Alexandra Swanson for helpful advice on spatial analyses, and Emilie Snell-Rood, David Stephens, Michael Wilson, and Lindsay Leverett for feedback on this manuscript. Permission to conduct this research was granted by the Guyana EPA (Permit No. 060214 BR 018 and 040717 BR 004) and the Guyana Protected Areas Commission. All research procedures were approved under UMN Institutional Animal Care and Use Protocol #1701-34456A.

## SUPPLEMENTAL FIGURES

**Figure S1.**
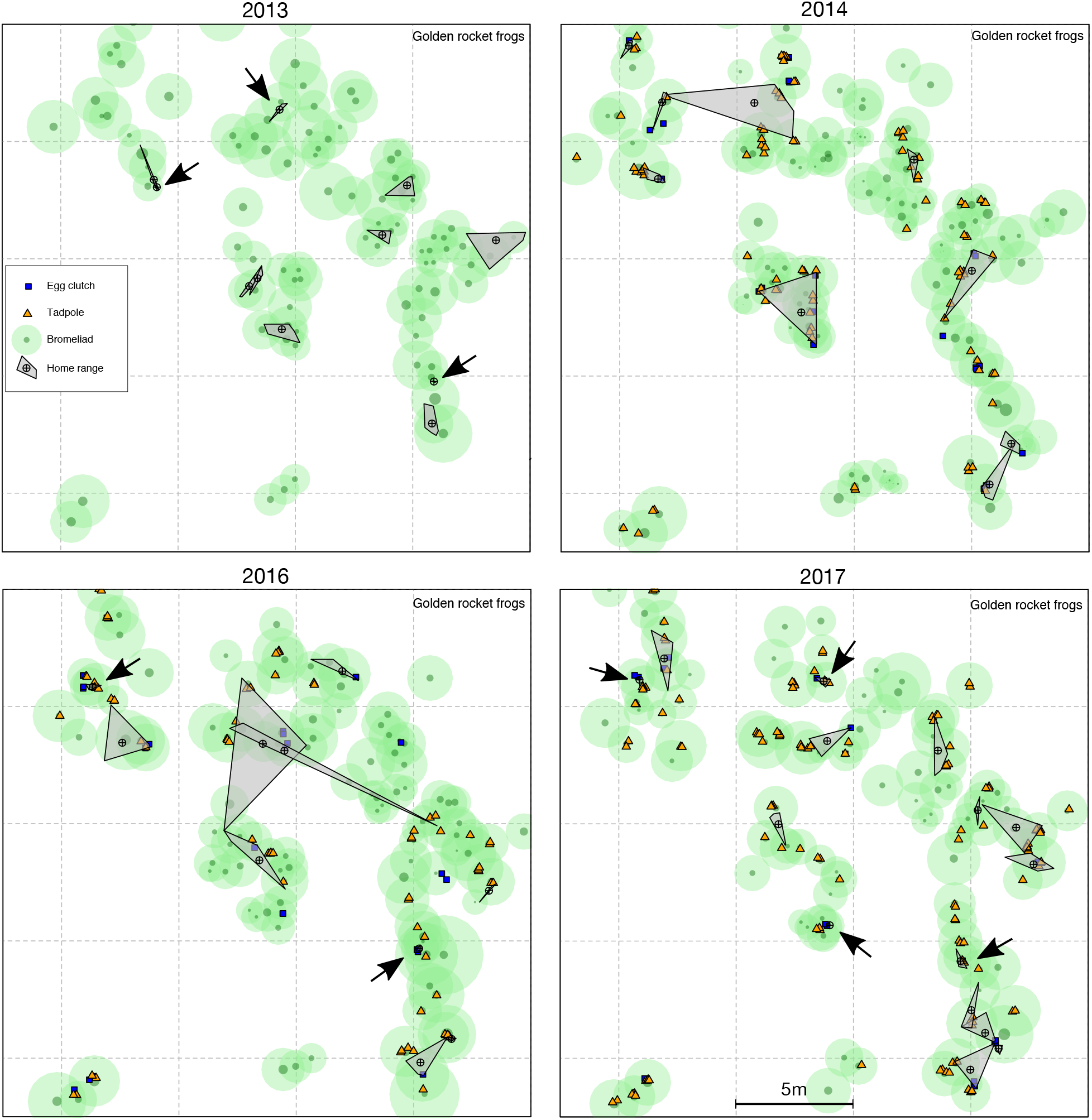
Maps of a site surveyed for golden rocket frogs over 5 years (the map for 2015 is shown in the main text). Filled grey polygons represent the home ranges of individual males, and black arrows point to males with particularly small home ranges. Light green circles represent the leaf diameters of bromeliads, while smaller dark green circles represent the phytotelmata diameter within bromeliads. Blue squares represent the locations of egg clutches and orange triangles represent the locations of tadpoles. Note: we did not map eggs or tadpoles in 2013.

**Figure S2.**
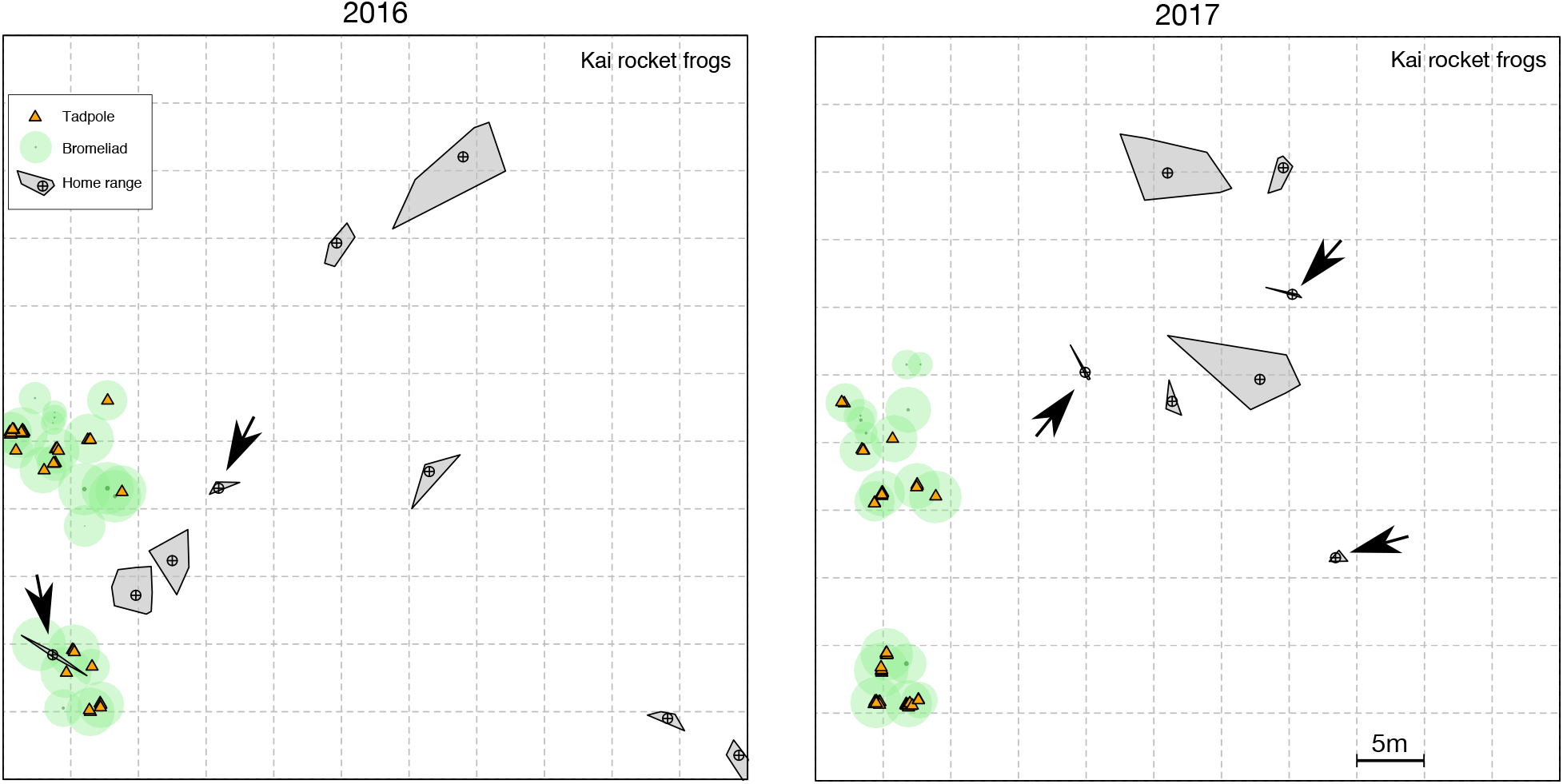
Maps of a site surveyed for male Kai rocket frogs over 3 years (the map for 2015 is shown in the main text). Filled grey polygons represent the home ranges of individual males, and black arrows point to males with particularly small home ranges. Light green circles represent the leaf diameters of bromeliads, while smaller dark green circles represent the phytotelmata diameter within bromeliads. Orange triangles represent the locations of tadpoles.

## REFERENCES

Bates D, Maechler M, Bolker B, Walker S. 2015. Fitting linear mixed-effects models using lme4. J Stat Softw. 67(1):1–48.

Bee MA. 2003. A test of the “dear enemy effect” in the strawberry dart-poison frog (*Dendrobates pumilio*). Behav Ecol Sociobiol. 54(6):601–610. doi:10.1007/s00265-003-0657-5.

Bee MA. 2016. Social recognition in anurans. In: Bee MA, Miller CT, editors. Psychological Mechanisms in Animal Communication. Springer. p. 169–221.

Bee MA, Gerhardt HC. 2002. Individual voice recognition in a territorial frog (*Rana catesbeiana*). Proc R Soc London B Biol Sci. 269(1499):1443–1448. doi:10.1098/rspb.2002.2041.

Bee MA, Reichert MS, Tumulty J. 2016. Assessment and recognition of competitive rivals in anuran amphibians. Adv Study Behav. 48:161–249. doi:10.1016/bs.asb.2016.01.001.

Bee MA, Schwartz JJ, Summers K. 2013. All’s well that begins Wells: celebrating 60 years of Animal Behaviour and 36 years of research on anuran social behaviour. Anim Behav. 85(1):5–18. doi:10.1016/j.anbehav.2012.10.031.

Beecher MD. 1991. Successes and failures of parent-offspring recognition in animals. In: Hepper PG, editor. Kin Recognition. Cambridge: Cambridge University Press. p. 94–124.

Beecher MD, Beecher IM, Lumpkins S. 1981. Parent-offspring recognition in bank swallows (*Riparia riparia*): I. Natural History. Anim Behav. 29:86–94.

Beletsky LD, Orians GH. 1989. Familiar neighbors enhance breeding success in birds. Proc Natl Acad Sci U S A. 86(20):7933–6.

Bergman TJ, Beehner JC. 2015. Measuring social complexity. Anim Behav. 103:203–209. doi:10.1016/j.anbehav.2015.02.018.

Bivand RS, Pebesma E, Gomez-Rubio V. 2013. Applied spatial data analysis with R. 2nd ed. New York: Springer.

Bivand RS, Rundel C. 2017. rgeos: Interface to Geometry Engine - Open Source (‘GEOS’).

Booksmythe I, Jennions MD, Backwell PRY. 2010. Investigating the ‘dear enemy’ phenomenon in the territory defence of the fiddler crab, *Uca mjoebergi*. Anim Behav. 79(2):419–423. doi:10.1016/j.anbehav.2009.11.020.

Bourne GR, Collins A, Holder A, McCarthy C. 2001. Vocal communication and reproductive behavior of the frog *Colostethus beebei* in Guyana. J Herpetol. 35(2):272–281.

Briefer E, Rybak F, Aubin T. 2008. When to be a dear enemy: flexible acoustic relationships of neighbouring skylarks, *Alauda arvensis*. Anim Behav. 76(4):1319–1325. doi:10.1016/j.anbehav.2008.06.017.

Brown JL, Morales V, Summers K. 2009. Home range size and location in relation to reproductive resources in poison frogs (Dendrobatidae): a Monte Carlo approach using GIS data. Anim Behav. 77(2):547–554. doi:10.1016/j.anbehav.2008.10.002.

Cheney DL, Seyfarth RM. 1999. Recognition of other individuals’ social relationships by female baboons. Anim Behav. 58(1):67–75. doi:10.1006/anbe.1999.1131.

Christensen C, Radford AN. 2018. Dear enemies or nasty neighbors? Causes and consequences of variation in the responses of group-living species to territorial intrusions. Behav Ecol. 29(5): 1004–1013. doi:10.1093/beheco/ary010.

Chuang M-F, Bee MA, Kam Y-C. 2013. Short amplexus duration in a territorial anuran: A possible adaptation in response to male-male competition. PLoS One. 8(12):e83116. doi:10.1371/journal.pone.0083116.

Chuang M-F, Kam Y-C, Bee MA. 2017. Territorial olive frogs display lower aggression towards neighbours than strangers based on individual vocal signatures. Anim Behav. 123:217–228. doi:10.1016/j.anbehav.2016.11.001.

Davis MS. 1987. Acoustically mediated neighbor recognition in the North American bullfrog, *Rana catesbeiana*. Behav Ecol Sociobiol. 21(3):185–190. doi:10.1007/BF00303209.

Donnelly MA. 1989. Demographic effects of reproductive resource supplementation in a territorial frog, *Dendrobates pumilio*. Ecol Monogr. 59(3):207–221.

Dyson ML, Reichert MS, Halliday TR. 2013. Contests in amphibians. In: Hardy ICW, Briffa M, editors. Animal Contests. Cambridge University Press. p. 228–257.

Fincke OM. 1992. Consequences of larval ecology for territoriality and reproductive success of a Neotropical damselfly. Ecology. 73(2):449–462.

Freeberg TM, Dunbar RIM, Ord TJ. 2012. Social complexity as a proximate and ultimate factor in communicative complexity. Philos Trans R Soc B Biol Sci. 367(1597):1785–801. doi:10.1098/rstb.2011.0213.

Gaulin SJC, Knight DH, Gaulin CK. 2018. Local variance in *Alouatta* group size and food availability on Barro Colorado Island. Biotropica. 12(2):137–143.

Gerhardt HC, Huber F. 2002. Acoustic Commmunication in Insects and Anurans. Chicago: University of Chicago Press.

Getty T. 1987. Dear enemies and the prisoner’s dilemma: why should territorial neighbors form defensive coalitions? Am Zool. 27(2):327–36.

Getty T. 1989. Are dear enemies in a war of attrition? Anim Behav. 37:337–9.

Grant T, Rada M, Anganoy-Criollo M, Batista A, Dias PH, Jeckel AM, Machado DJ, Rueda-Almonacid JV. 2017. Phylogenetic systematics of dart-poison frogs and their relatives revisited (Anura: Dendrobatoidea). South Am J Herpetol. 12(s1):S1–S90. doi:10.2994/SAJH-D-17-00017.1.

Hatchwell BJ, Komdeur J. 2000. Ecological constraints, life history traits and the evolution of cooperative breeding. Anim Behav. 59(6):1079–1086. doi:10.1006/anbe.2000.1394.

Howard RD. 1978. The evolution of mating strategies in bullfrogs, *Rana catesbeiana*. Evolution. 32(4):850–871.

Humfeld SC, Marshall VT, Bee MA. 2009. Context-dependent plasticity of aggressive signalling in a dynamic social environment. Anim Behav. 78(4):915–924. doi:10.1016/j.anbehav.2009.06.028.

Hyman J. 2005. Seasonal variation in response to neighbors and strangers by a territorial songbird. Ethology. 111(10):951–961. doi:10.1111/j.1439-0310.2005.01104.x.

Jaeger RG. 1981. Dear enemy recognition and the costs of aggression between salamanders. Am Nat. 117(6):962–974.

Kok PJ, Kalamandeen M. 2008. Introduction to the taxonomy of the amphibians of Kaieteur National Park, Guyana. ABC Taxa. 5.

Kok PJ, Sambhu H, Roopsind I, Lenglet GL, Bourne GR. 2006. A new species of *Colostethus* (Anura: Dendrobatidae) with maternal care from Kaieteur National Park, Guyana. Zootaxa. 1238(May):35–61.

Kumar S, Stecher G, Suleski M, Hedges SB. 2017. TimeTree: a resource for timelines, timetrees, and divergence times. Mol Biol Evol. 34(7):1812–1819. doi:10.1093/molbev/msx116.

Marshall VT, Humfeld SC, Bee MA. 2003. Plasticity of aggressive signalling and its evolution in male spring peepers, *Pseudacris crucifer*. Anim Behav. 65:1223–1234. doi:10.1006/anbe.2003.2134.

Medvin MB, Beecher MD. 1986. Parent-offspring recognition in the barn swallow (*Hirundo rustica*). Anim Behav. 34:1627–1639.

Miller DB. 1979. The acoustic basis of mate recognition by female Zebra finches (*Taeniopygia guttata*). Anim Behav. 27:376–380. doi:10.1016/0003-3472(79)90172-6.

Müller CA, Manser MB. 2007. “Nasty neighbours” rather than “dear enemies” in a social carnivore. Proc R Soc B. 274(1612):959–965. doi:10.1098/rspb.2006.0222.

Ostfeld RS. 1985. Limiting resources and territoriality in microtine rodents. Am Nat. 126(1):1–15. doi:10.1086/521238.

Pettitt BA, Bourne GR, Bee MA. 2012. Quantitative acoustic analysis of the vocal repertoire of the golden rocket frog (*Anomaloglossus beebei*). J Acoust Soc Am. 131(6):4811–20. doi:10.1121/1.4714769.

Pettitt BA, Bourne GR, Bee MA. 2013. Advertisement call variation in the golden rocket frog (*Anomaloglossus beebei*): evidence for individual distinctiveness. Ethology. 119:244–256.

Pettitt BA, Bourne GR, Bee MA. 2018. Predictors and benefits of microhabitat selection for offspring deposition in golden rocket frogs. Biotropica. 50(6):919–928. doi:10.1111/btp.12609.

Pettitt BA, Bourne GR, Bee MA. 2019. Females prefer the calls of better fathers in a Neotropical frog with biparental care. Behav Ecol. 31(1):152–163. doi:10.1093/beheco/arz172.

Poelman EH, Dicke M. 2008. Space use of Amazonian poison frogs: testing the reproductive resource defense hypothesis. J Herpetol. 42(2):270–278. doi:10.1670/07-1031.1.

Pröhl H. 1997. Territorial behaviour of the strawberry poison-dart frog, *Dendrobates pumilio*. Amphibia-Reptilia. 18:442–446. doi:10.1017/CBO9781107415324.004.

Pröhl H. 2005. Territorial behavior in dendrobatid frogs. J Herpetol. 39(3):354–365.

Pröhl H, Berke O. 2001. Spatial distributions of male and female strawberry poison frogs and their relation to female reproductive resources. Oecologia. 129(4):534–542. doi:10.1007/S004420100751.

Qualls CP, Jaeger RG. 1991. Dear enemy recognition in *Anolis carolinensis*. J Herpetol. 25(3):361–363.

R Core Team. 2017. R: A language and environment for statistical computing.

Reeve HK. 1989. The evolution of conspecific acceptance thresholds. Am Nat. 133(3):407–435.

Reichert MS. 2010. Aggressive thresholds in *Dendropsophus ebraccatus:* Habituation and sensitization to different call types. Behav Ecol Sociobiol. 64(4):529–539. doi:10.1007/s00265-009-0868-5.

Ringler M, Ringler E, Magaña Mendoza D, Hödl W. 2011. Intrusion experiments to measure territory size: development of the method, tests through simulations, and application in the frog *Allobates femoralis*. PLoS One. 6(10):e25844. doi:10.1371/journal.pone.0025844.

Ringler M, Ursprung E, Hödl W. 2009. Site fidelity and patterns of short- and long-term movement in the brilliant-thighed poison frog *Allobates femoralis* (Aromobatidae). Behav Ecol Sociobiol. 63(9):1281–1293. doi:10.1007/s00265-009-0793-7.

Roithmair ME. 1992. Territoriality and male mating success in the dart-poison frog, *Epipedobates femoralis* (Dendrobatidae, Anura). Ethology. 92(4):331–343.

Rose GJ, Brenowitz EA. 1991. Aggressive thresholds of male Pacific treefrogs for advertisement calls vary with amplitude of neighbors calls. Ethology. 89:244–252.

Rubenstein DR, Lovette IJ. 2007. Temporal environmental variability drives the evolution of cooperative breeding in birds. Curr Biol. 17(16):1414–1419. doi:10.1016/j.cub.2007.07.032.

Sheehan MJ, Tibbetts EA. 2009. Evolution of identity signals: frequency-dependent benefits of distinctive phenotypes used for individual recognition. Evolution. 63(12):3106–13. doi:10.1111/j.1558-5646.2009.00833.x.

Sheehan MJ, Tibbetts EA. 2010. Selection for individual recognition and the evolution of polymorphic identity signals in *Polistes* paper wasps. J Evol Biol. 23(3):570–7. doi:10.1111/j.1420-9101.2009.01923.x.

Sherman PW, Reeve HK, Pfennig DW. 1997. Recognition systems. In: Krebs JR, Davies NB, editors. Behavioral ecology: An evolutionary approach. 4th ed. Oxford: Blackwell Science Ltd. p. 69–96.

Summers K. 1992. Mating strategies in two species of dart-poison frogs: a comparative study. Anim Behav. 43:907–919.

Summers K. 2000. Mating and aggressive behaviour in dendrobatid frogs from Corcovado National Park, Costa Rica: a comparative study. Behaviour. 137:7–24.

Summers K, Tumulty J. 2013. Parental care, sexual selection, and mating systems in neotropical poison frogs. In: Macedo RH, Machado G, editors. Sexual Selection: Perspectives and Models from the Neotropics. Waltham, MA: Elsevier Academic Press.

Temeles E. 1994. The role of neighbours in territorial systems: when are they “dear enemies”? Anim Behav. 47:339–350.

Tibbetts EA, Dale J. 2007. Individual recognition: it is good to be different. Trends Ecol Evol. 22(10):529–37. doi:10.1016/j.tree.2007.09.001.

Tumulty JP. 2018a. Dear Enemy Effect. In: Vonk J, Shackelford T, editors. Encyclopedia of Animal Cognition and Behavior. Cham: Springer International Publishing. p. 1–4.

Tumulty JP. 2018b. The evolution and mechanisms of social recognition in territorial frogs [dissertation]. University of Minnesota.

Tumulty JP, Pašukonis A, Forester JD, Ringler M, Hödl W, Bee MA. 2018. Brilliant-thighed poison frogs do not use acoustic identity information to treat territorial neighbours as dear enemies. Anim Behav. 141:203–220.

Tumulty JP, Sheehan MJ. 2020. What drives diversity in social recognition mechanisms? Front Ecol Evol. 7(517). doi:10.3389/fevo.2019.00517.

Ursprung E, Ringler M, Jehle R, Hödl W. 2011. Strong male/male competition allows for nonchoosy females: high levels of polygynandry in a territorial frog with paternal care. Mol Ecol. 20(8):1759–1771. doi:10.1111/j.1365-294X.2011.05056.x.

Vacher JP, Kok PJR, Rodrigues MT, Lima JD, Lorenzini A, Martinez Q, Fallet M, Courtois EA, Blanc M, Gaucher P, et al. 2017. Cryptic diversity in Amazonian frogs: Integrative taxonomy of the genus *Anomaloglossus* (Amphibia: Anura: Aromobatidae) reveals a unique case of diversification within the Guiana Shield. Mol Phylogenet Evol. 112:158–173. doi:10.1016/j.ympev.2017.04.017.

Wells KD. 1977. The social behaviour of anuran amphibians. Anim Behav. 25:666–693.

Wells KD. 2007. The Ecology and Behavior of Amphibians. Chicago: University of Chicago Press.

Werner P, Elle O, Schulte LM, Lötters S. 2010. Home range behaviour in male and female poison frogs in Amazonian Peru (Dendrobatidae: *Ranitomeya reticulata*). J Nat Hist. 45(1-2):15–27. doi:10.1080/00222933.2010.502257.

Wiley RH. 2013a. Specificity and multiplicity in the recognition of individuals: implications for the evolution of social behaviour. Biol Rev. 88:179–95. doi:10.1111/j.1469-185X.2012.00246.x.

Wiley RH. 2013b. A receiver-signaler equilibrium in the evolution of communication in noise. Behaviour. 150(9-10):957–993. doi:10.1163/1568539X-00003063.

Wilson EO. 1975. Sociobiology: The New Synthesis. Cambridge, MA: Harvard University Press.

Ydenberg RC, Giraldeau L-A, Falls JB. 1988. Neighbours, strangers, and the asymmetric war of attrition. Anim Behav. 36:343–347.

